# In utero human intestine contains maternally derived bacterial metabolites

**DOI:** 10.1101/2024.08.20.608888

**Authors:** Wenjia Wang, Weihong Gu, Ron Schweitzer, Omry Koren, Soliman Khatib, George Tseng, Liza Konnikova

## Abstract

Understanding when host-microbiome interactions are first established is crucial for comprehending normal development and identifying disease prevention strategies. Furthermore, bacterially derived metabolites play critical roles in shaping the intestinal immune system. Recent studies have demonstrated that memory T cells infiltrate human intestinal tissue early in the second trimester, suggesting that intestinal immune education begins in utero. Our previous study reported a unique fetal intestinal metabolomic profile with an abundance of several bacterially derived metabolites and aryl hydrocarbon receptor (AHR) ligands implicated in mucosal immune regulation. To follow up on this work, in the current study, we demonstrate that a number of microbial byproducts present in fetal intestines in utero are maternally derived and vertically transmitted to the fetus. Notably, these bacterially derived metabolites, particularly short chain fatty acids and secondary bile acids, are likely biologically active and functional in regulating the fetal immune system and preparing the gastrointestinal tract for postnatal microbial encounters, as the transcripts for their various receptors and carrier proteins are present in second trimester intestinal tissue through single-cell transcriptomic data.

## Introduction

Establishing and maintaining a “healthy” intestinal microbiome is critical to the overall health of an individual. This is partly facilitated by bacterially derived metabolites that are important mediators of intestinal health and regulators of intestinal mucosal immunity. Furthermore, disruption in this homeostasis can lead to a myriad of diseases (1, 2). Several groups of metabolites have been identified that are particularly important to the gut homeostasis including short-chain fatty acids (SCFA), and secondary bile acids, among others (3). Understanding how and when the dialogue between the intestinal tract (host) and the bacterial metabolites is established is important to designing new strategies in preventing disease and improving health. There has recently been a lot of attention on the development of the microbiome over the first three years of life (or the first 1000 days) as being a critical window to modulate the development and adaptation of the immune system and overall health (4). With the detection of the memory T cells within the human and non-human primate fetal intestines (5–8), cord blood (9), and placenta (10), among other tissues, scientists are appreciating that education of the intestinal immune system may be ongoing in utero and the critical window of shaping the intestinal homeostasis potentially starts prior to delivery (11). Yet almost nothing is known about the antigens present in utero or the metabolites modulating the intestinal homeostasis.

Several studies have reported that the placenta lacks a microbiome (12–16) reviewed in (17), and others report minimal bacterial colonization in fetal meconium (18), as well as placental and endometrial samples (19–24). Furthermore, research conducted by Lauder et al. (13), involving a large cohort, found no discernible difference between placental samples and kit samples (contamination introduced during DNA purification). However, pioneering work from the MacPherson’s group demonstrated that metabolites from the maternal intestinal microbiome can be detected in the murine fetal intestine and alter the development of the fetal mucosal immune system (25). Building on this, our previous study found a unique fetal intestinal metabolomic profile with an abundance of bacterially derived metabolites and aryl hydrocarbon receptor (AHR) ligands implicated in mucosal immune regulation (26).

In the current study, we hypothesized that microbial byproducts detected in fetal intestines in utero are primarily derived from the maternal microbiota and play a role in preparing the intestinal immune system for ex-utero life. To study this, we assembled a cohort of pregnancy-matched fetal organs including the fetal intestine, the fetal meconium, the fetal placental villi, to the maternal decidua. We found that some of the microbial byproducts or metabolites present in fetal intestines in utero were maternally derived and vertically transmitted to the fetus, while some were found to be in a steady state. Importantly, these bacterially derived metabolites are likely biologically active and functional in regulating the fetal immune system and prepare the gastrointestinal tract for postnatal microbial encounters as we were able to detect their receptors and carrier proteins in single cell transcriptomic data of the human fetal small intestine.

## Results

To investigate the source of *in utero* intestinal metabolome, we performed untargeted metabolomic analysis, including human and bacterially derived metabolites (manually curated to be fully bacterially derived or require partial conversion by the bacteria), using in house pipeline established by the Khatib lab (26) on 49 tissue samples from 24 subjects (with gestational age ranging from 14 weeks to 23 weeks): 8 maternal decidua samples, 11 fetal placental villi (PV) samples, 11 fetal gastrointestinal (GI) (fetal small intestine (SI) and large intestine (LI)) samples, and 19 fetal meconium samples (**Table 1**).

**Table 1.**
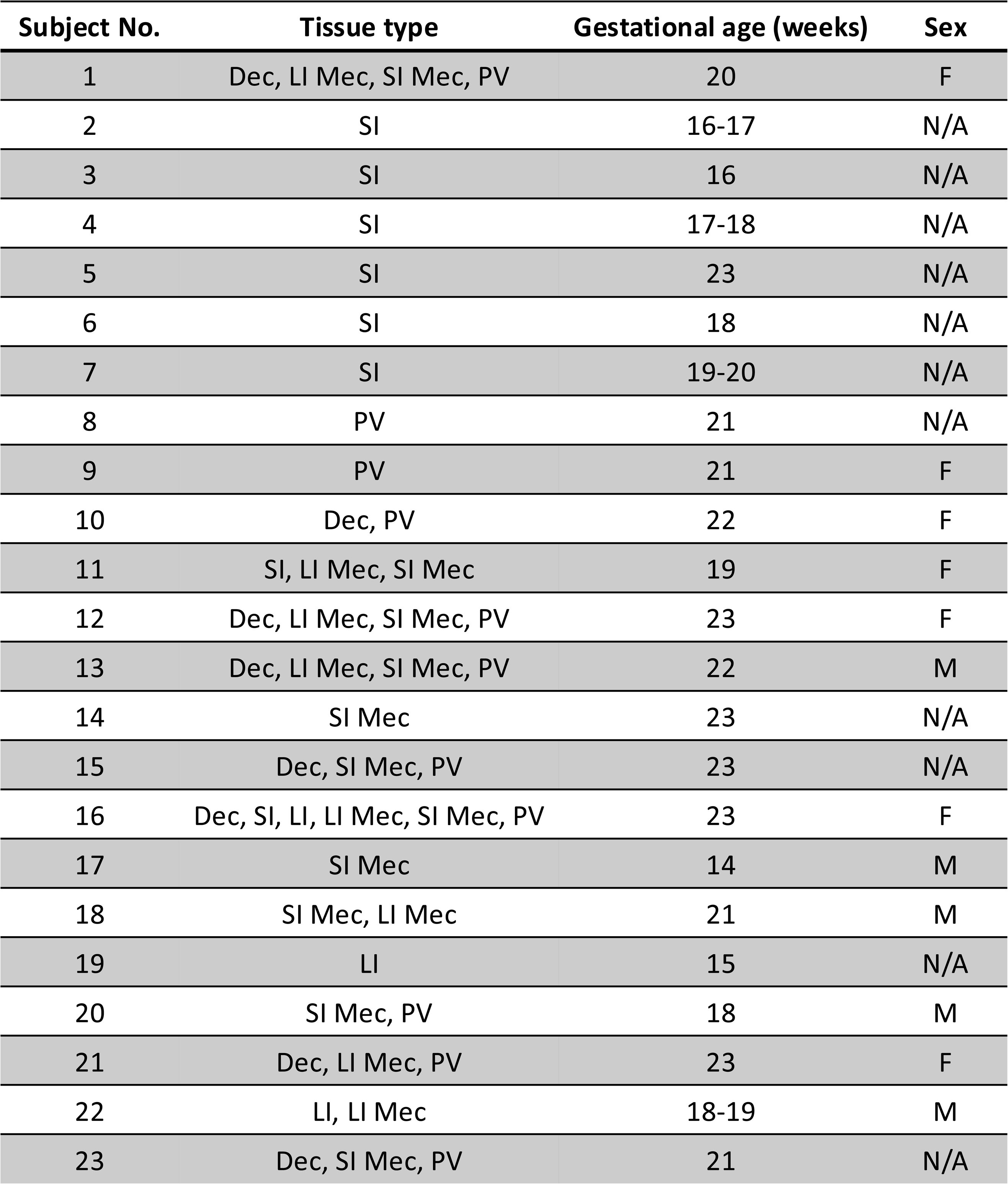
Sample demographics. Demographic information for all samples used. N/A, not available; SI, small intestine; LI, large intestine; SI Mec, small intestine meconium; LI Mec, large intestine meconium; F, female; M, male.

A total of 2521 metabolites were identified in all the 49 samples (**Supplementary Table S1**). t-SNE visualization based on the 2521 analyzed metabolites showed that samples of GI, meconium, decidua and PV were well separated, although some PV samples mixed with decidua samples possibly due to incomplete tissue separation. The separation indicated that even though most metabolites were detected across all samples, they were differentially abundant based on tissue source (**Figure 1A**). The top 20 GI enriched metabolites had differential abundances between GI and the other tissue groups are shown in **Figure 1B**. To explore tissue-specific individual metabolite signatures in the decidua, PV and meconium groups, we performed differential metabolite abundance analysis between GI and each of the other groups (see details in **Supplementary Table S2**). We detected a large proportion of metabolites whose abundance differed between tissue: GI vs. decidua =326, 13% (**Figure 1C**); GI vs. meconium= 318, 13% (**Figure 1D**); and GI vs. PV = 324, 13% (**Figure 1E**). Specifically, among the 326 differentially abundant metabolites between GI and decidua tissue, more than half (195 metabolites) were enriched for in the GI tissue, two of which were bacterial metabolites (Coprocholic acid and Glycodeoxycholic acid, secondary bile acids). For the 131 metabolites significantly enriched in decidua, five were bacterial metabolites (Pyridoxine, Methylhippuricacid, Hippurate, 2-Hydroxyhippuric acid, and Indoxyl sulfate). In the comparison between GI and meconium, 164 metabolites had higher abundance in GI tissue including three bacterial metabolites (N-Acetyl-alpha-D-glucosamine1-phosphate, Kynurenine, and Phosphopantothenic acid). Interestingly, two bacterial metabolites (benzoate and Albaflavenol) were enriched for in meconium. At the same time, there were six xenobiotics enriched for in the meconium, suggesting that even maternally ingested compounds can be concentrated in the meconium. Comparisons between the GI and PV tissue identified 235 metabolites that were enriched for in the GI tissue, with two bacterial metabolites (Coprocholic acid and Glycodeoxycholic acid), four primary bile acids (Glycocholic acid, 7-Sulfocholic acid, Taurocholic acid and Taurochenodeoxycholic acid), one aromatic acid (D-(+)-Tryptophan) and two xenobiotics (Adaprolol and Elacytarabine). Among the 89 metabolites with higher abidance in the PV tissue, there were two bacterial metabolites (Methyl hippuric acid and Indoxyl sulfate), and two xenobiotics (Penicillin-G and a THC derivative).

**Figure 1.**
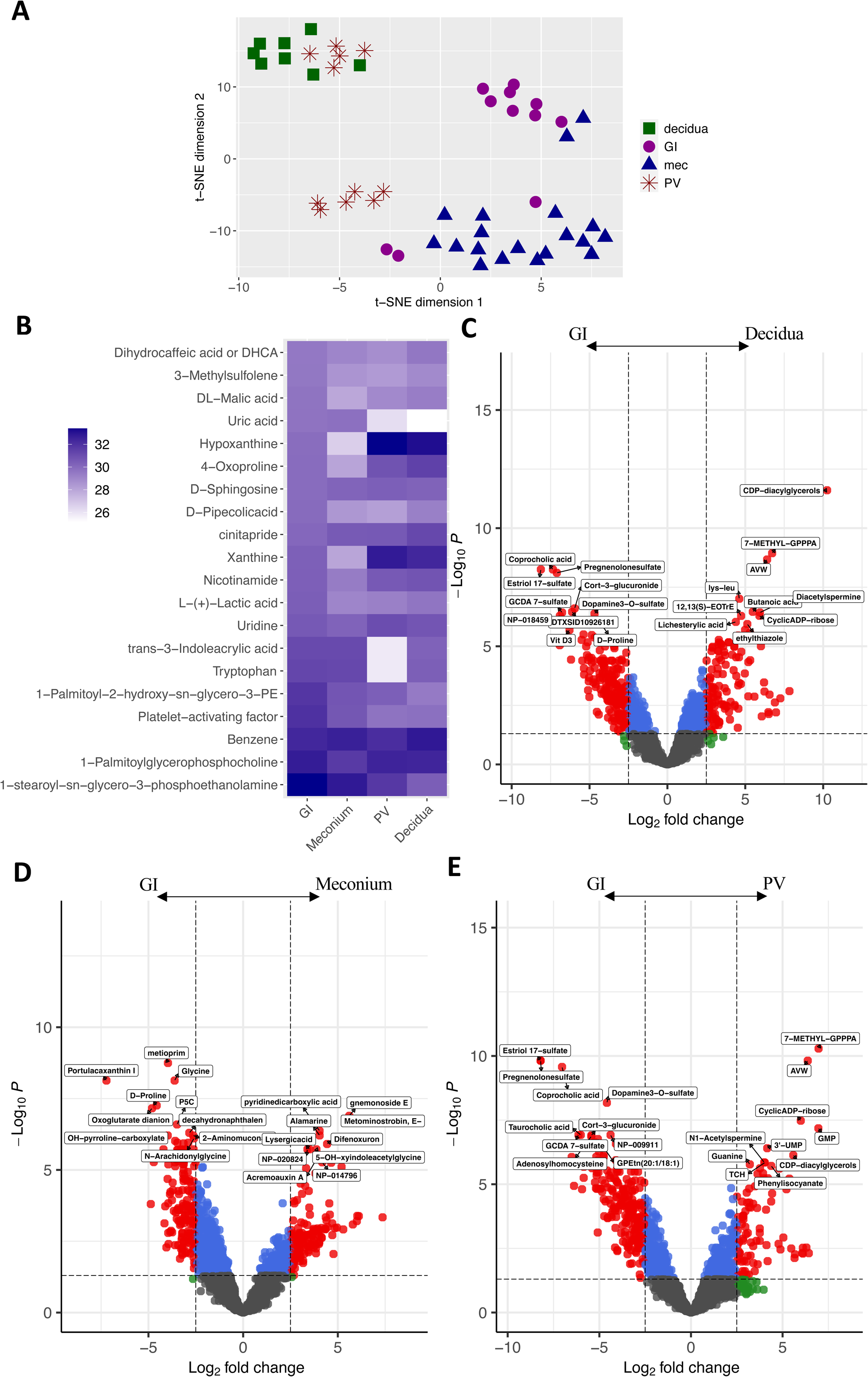
Sample separation and differential expression of individual metabolites. (A) t-distributed stochastic neighbor embedding (t-SNE) plot using all metabolites. (B) Heatmap showing normalized abundance of the top 20 expressed metabolites in the fetal GI samples (SI and LI), as well as their abundance in the meconium (SI Mec and LI Mec), PV, and maternal decidua samples. (C), (D) and (E) Volcano plots of differentially abundant metabolites between the GI and decidua groups, GI and meconium groups, and GI and PV groups respectively. Top ten most abundant metabolites are labeled with the metabolite name.

To understand which pathways are differentially regulate in the different tissue, we conducted Ingenuity Pathway Analysis (IPA) (see details in **Supplementary Table S3**). Of the pathways that were significantly altered between the decidua and GI samples, majority were more activated in the decidua, including: the transport of vitamins and steroid metabolism consistent with known functions of the placenta, while tryptophan catabolism and bile acid transport were more active in the GI track **(Figure 2A**). Very few pathways were more active in the meconium. Interestingly, pathways classically associate with post-natal intestinal epithelial function, were already enriched for prenatally including upregulation of many metabolism and transport pathways, prostaglandin synthesis, and neurotransmitter release (**Figure 2B**). Our group had previously discovered that human fetal intestinal cells can produce insulin and respond to glucose concentrations (27), in support of this, the current data identified that insulin secretion was upregulated in the fetal GI tract (**Figure 2B**). Bile acid transport pathways were also upregulated in the GI samples compared to PV samples (**Figure 2C**).

**Figure 2.**
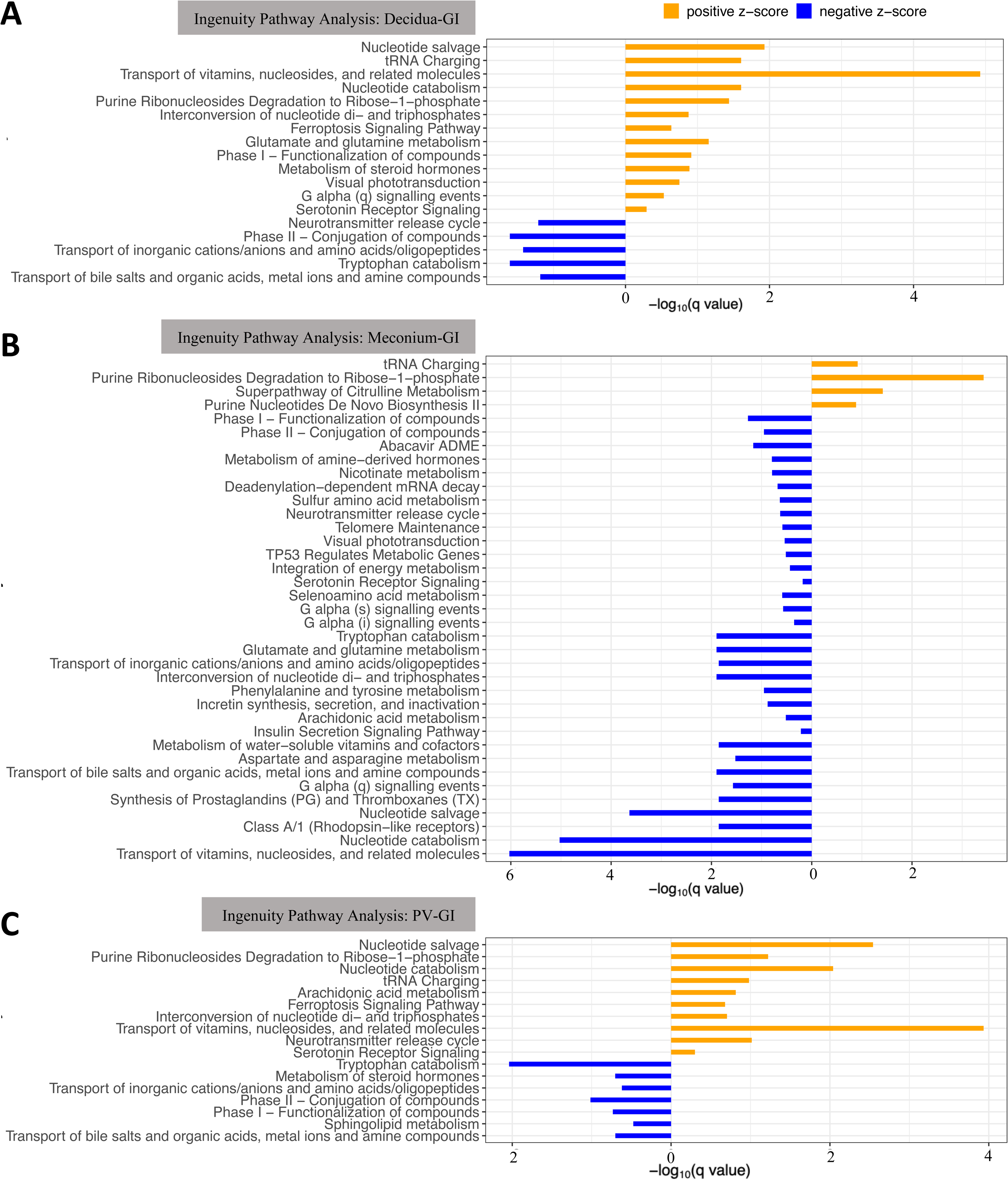
Pathway enrichment. (A), (B) and (C) Integrated pathway analysis for differentially altered pathways between GI and decidua groups, between GI and meconium groups, and between GI and PV groups. The length of the bar is proportional to the q value. Pathways with positive values on the x axis (orange bar) are those enriched for in the decidua, meconium, and PV respectively. Those with negative values (blue bars) are pathways enriched for in the GI samples.

To understand the source of the bacterial metabolites present within the fetal intestine and determine if they are vertically transmitted from the maternal microbiota, we performed correlation analysis between the fetal intestinal tissue and the decidual tissue (maternal origin) and fetal intestine and the meconium (local production, **Figure 3A**). Among the 2521 metabolites, we identified and selected 41 microbially derived or bacteria associated metabolites of interest based on a literature search, including 5 secondary bile acids, 3 short chain fatty acids (SCFA) and 3 aromatic lactic acids. In addition, we identified 47 xenobiotics, metabolites that cannot be produced in the human host, and 13 metabolites enriched for in the fetal tissue (**Supplementary Table S4**). All bacterial metabolites were present in the GI tissue, but the four tissues had different signatures of these metabolites (**Figure 3B**). To understand the source of the metabolites better, we used the matched tissue from one subject that had all tissues in the study from the same individual to perform correlation analysis. As expected, the SI and LI and the decidua and PV samples that come from adjacent tissue had the highest correlation of all samples, validating our analysis. Interestingly, the paired fetal GI and decidua samples had substantially higher positive correlation than the paired GI and meconium samples in terms of the abundance of the 41 microbial metabolites: p^Dec_SI^ = 0.88, p^Dec_LI^ = 0.88 while p^MecSI_SI^ = 0.62, p^MecLI_SI^ = 0.58, p^MecSI_LI^ = 0.66, p^MecLI_LI^ = 0.61 (**Figure 3C**), suggesting that the microbial metabolites detected in fetal samples were more likely to be vertically transmitted from maternal microbiota. To determine if this positive correlation also held true for all the samples in the study, we performed the two correlative analysis (GI/decidua and GI/meconium) using all samples combined for xenobiotics (expecting the ratio of GI/decidua to be more positively correlated than GI/mec as these can only come from maternal circulation), fetally derived metabolites (expecting the ratio of GI/decidua to be less than GI/meconium as these are locally produced in the fetus) and of bacteria associated metabolites. As expected, there was a significantly higher correlation based on the 47 xenobiotics between GI samples and decidua samples compared to the corresponding correlation between GI samples and meconium samples (**Figure 3D**). There was also a lower correlation based on the 13 fetal derived metabolites between GI samples and decidua samples compared to the corresponding correlation between GI samples and meconium samples (**Figure 3D**). Consistent with data from **Figure 3C** from an individual subject, we also observed that bacterially derived metabolites had a higher correlation between GI samples and decidua samples compared to the corresponding correlation between GI samples and meconium samples (**Figure 3D**).

**Figure 3.**
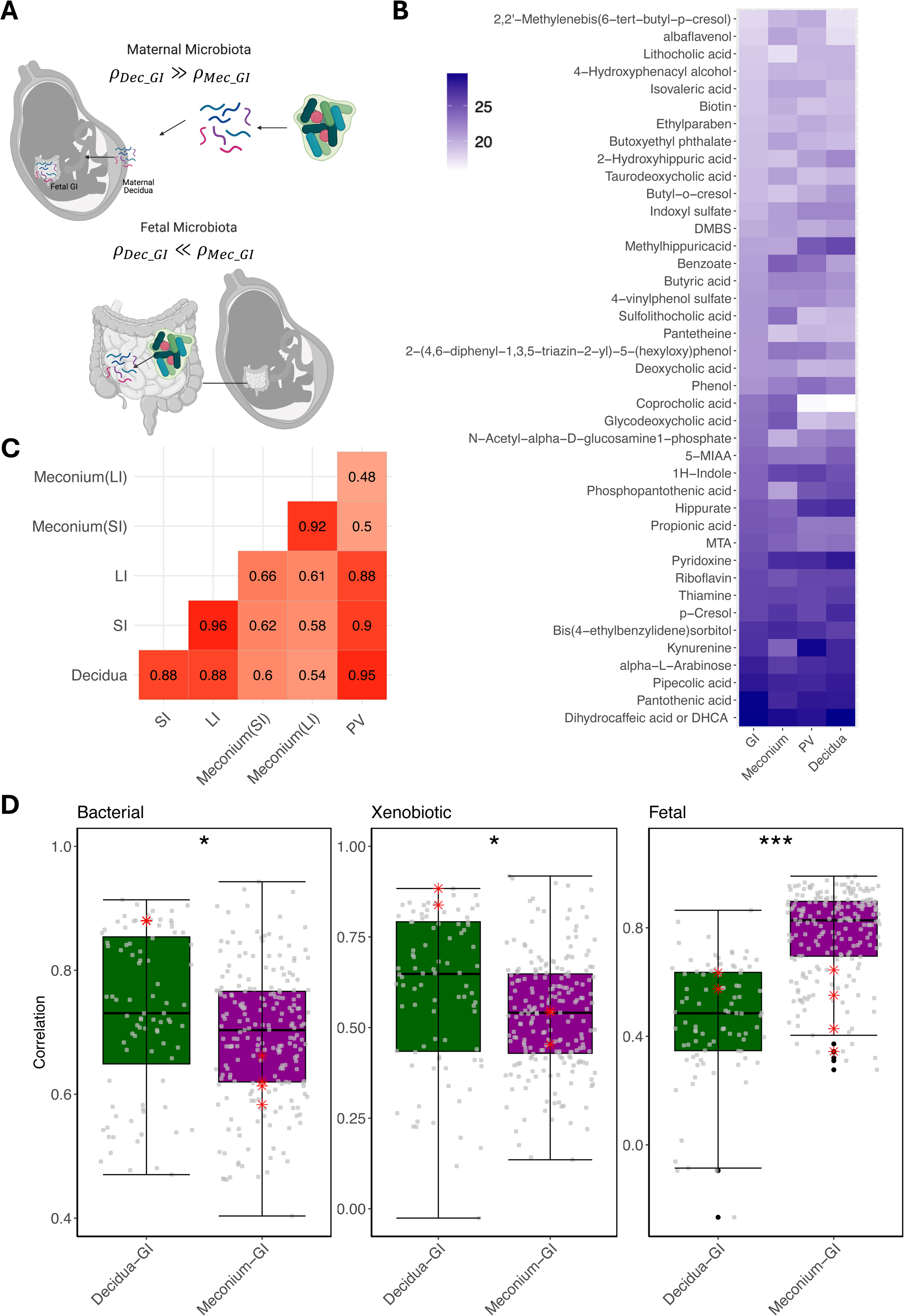
Correlation between tissue groups of microbial metabolites, xenobiotics, and fetal metabolites. (A) Schematic of how the correlation between tissue would identify the source of bacterial metabolites. (B) Heatmap showing normalized abundance of the 41 microbial associated metabolites across sample types. (C) Pairwise correlation matrix of the 41 microbial metabolites between paired tissue samples from subject 1146. (D) Boxplots visualizing the correlations between GI and decidua groups, and between GI and meconium groups based on 41 microbial metabolites, 47 xenobiotics, and 13 fetal-derived metabolites respectively from **Supplementary Table** S4. Red asterisk points represent the pairwise correlations between tissue samples from subject 1146. *P < 0.05, ***P<0.001.

Overall, the abundance of the 41 microbial metabolites identified in our dataset, demonstrated a high correlation between GI and decidua samples. To determine if there was variability by metabolite, we determined the abundance enrichment across the tissues for each metabolite. In our analysis, the primary bile acids that are produced in the fetal liver and then absorbed by the small intestine, as expected, were significantly enriched for in fetal meconium but deprived in maternal decidua except for muricholic acid whose abundance was similar across tissues (**Figure 4A**). Upon crossing the intestinal lumen, secondary bile acids are produced by microbiota mediated dehydroxylation or deconjugation of the bile acids (28). We were able to identify five secondary bile acids present in our dataset. Lithocholic acid was enhanced for in the decidua, while deoxycholic acid was present in similar abundance across all the tissues. Surprisingly, our analysis identified three secondary bile acids: Glycodeoxycholic acid, Sulfolithocholic acid, and Taurodeoxycholic acid whose abundance was significantly high in meconium samples.

**Figure 4.**
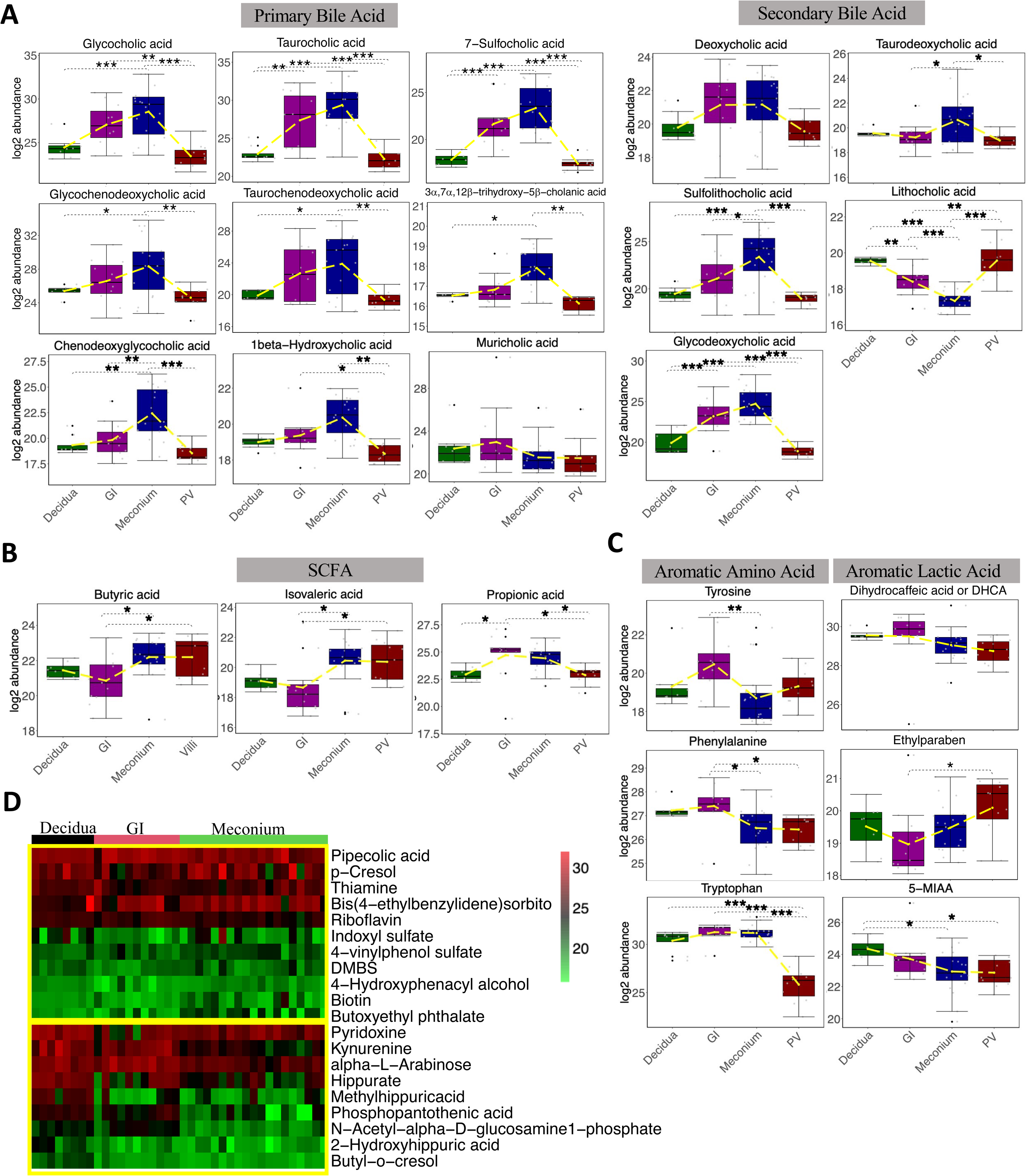
ANOVA analysis of SCFA, bile acids and aromatic acids across tissues. (A-C) Boxplots of individual metabolite’s abundance for primary and secondary bile acids, SCFA, and aromatic amino acids and aromatic acids. *P < 0.05, **P<0.01, ***P<0.001. (D) Heatmap showing that microbial metabolites without significant difference across tissue groups (upper panel) and microbial metabolites significantly enriched in decidua samples (lower panel).

We then determine if short chain fatty acids (SCFA), important source of nutrition for intestinal epithelial cells and modulators of immunity, that are usually locally produced within the intestinal lumen, were present in the fetal intestine. Three SCFA were detected in our dataset, butyric acid and isovaleric acid were significantly reduced in the GI tissue and enhanced in the meconium and the PV (**Figure 4B**). While proprionic acid was enriched for in the GI samples above all other tissue (**Figure 4B**).

Another important group of bacterially derived metabolites are aromatic lactic acids. Breastfeeding has been reported to promote Bifidobacterium species converting aromatic amino acids tyrosine, phenylalanine, and tryptophan into their respective aromatic lactic acids dihydrocaffeic acid or DHCA, ethylparaben, and 5-MIAA (5-Methoxyindoleacetate), that are biologically active in the intestine and associated with anti-inflammatory properties (29, 30). We additionally explored if these metabolites are present in fetal tissue. The three aromatic amino acids were similarly abundant in the decidua and the GI tract with tryptophan levels being reduced in the meconium (**Figure 4C**). All three of the aromatic lactic acids were present in fetal tissue, where 5-MIAA levels were lowest in the meconium and levels of DHCA and ethylparaben were similarly abundant between the decidua, GI and meconium samples (**Figure 4C**).

Finally, we evaluated the remaining bacterially associated metabolites present in fetal tissue. Nine of these microbial metabolites (Pyridoxine, N-Acetyl-alpha-D-glucosamine1-phosphate, alpha-L-Arabinose, Kynurenine, Methylhippuricacid, Phosphopantothenic acid, Hippurate, Butyl-o-cresol, and 2-Hydroxyhippuric acid) were significantly enriched for in decidua samples, while 11 microbial metabolites (Riboflavin, Pipecolic acid, 4-Hydroxyphenacyl alcohol, 4-vinylphenol sulfate, DMBS, Biotin, Butoxyethyl phthalate, Bis(4-ethylbenzylidene)sorbitol, Thiamine, p-Cresol, and Indoxyl sulfate) were found in similarly abundance across all tissue samples (**Figure 4D**).

To ensure that metabolites were identified correctly, a number of these were validated with known standards. These included: butyric acid, deoxycholic acid, pantetheine, p-Cresol, taurochenodeoxycholic acid, hippurate, benzoate, and taurocholic acid (**Supplementary Figure S1A-B**). Due to sample limitation where no additional sample was available, metabolites were validated by standards however the exact concentrations of the samples could not be calculated. Where additional samples were available, metabolites were validated by standards and exact concentrations were calculated. For deoxycholic acid, both methods were used. The results demonstrated consistent trends between the targeted and untargeted analysis among the four tissue groups for each metabolite except hippurate where quantitively analysis didn’t identify any difference in abundance between groups.

Intestinal bile acids have been shown to alter abundance and type of mucosal regulatory T cells. Interestingly, T cells begin to populate the small intestine in humans early on in the second trimester (31), suggesting that the differential presence of various bile acids across gestational ages may be important in establishing/regulating mucosal immunity. Similarly, SCFA have also been shown to play a role in mucosal immunity, particularly T cell homeostasis (32). Here we explored the association between the abundance of the identified nine primary bile acids, five secondary bile acids, and three SCFAs with gestational age within tissue groups. Three primary bile acids: muricholic acid, 1beta-Hydroxycholic acid, and 3a,7a,12b-trihydroxy-5b-cholanic acid had a significant positive association with advancing gestational age in the GI tissue, indicative of their increased synthesis in the fetus with advancing gestational age (**Table 2**). In contrast, although most of the bile acids and SCFAs did not have any significant association with gestational age (**Supplementary Table S5**), we found that two secondary bile acids, deoxycholic acid and glycodeoxycholic acid, and one SCFA, proprionic acid, had a significant negative correlation with advancing gestational age in the fetal intestine (**Table 2**). The negative correlation suggests these compounds decrease in the GI tract with gestational age, highlighting that they are likely coming from maternal circulation rather than local production.

**Table 2.** Correlation coefficients between the abundance of individual metabolites and the gestational age within each tissue group. Only show the five bile acids and one SCFA (Propionic acid) with significant age-association. The rows are the correlation coefficient within each tissue group separately. The q value for the correlation coefficient shows in parathesis *P < 0.05, **P<0.01, ***P<0.001.

| | Deoxycholic acid | Glycodeoxycholic acid | Propionic acid | Muricholic acid | 1beta-Hydroxycholic acid | 3α,7α,12β-trihydroxy-5β-cholanolic acid |
| --- | --- | --- | --- | --- | --- | --- |
| Decidua | -0.33<br>(0.94) | -0.38<br>(0.94) | -0.15<br>(0.94) | 0.37<br>(0.94) | 0.26<br>(0.94) | 0.09<br>(0.94) |
| GI | -0.29***<br>($<0.001$) | -0.19*<br>(0.05) | -0.25***<br>($<0.001$) | 0.45***<br>($<0.001$) | 0.35***<br>($<0.001$) | 0.26***<br>($<0.001$) |
| Meconium | 0.28<br>(0.43) | 0.29<br>(0.43) | 0.07<br>(0.76) | -0.24<br>(0.43) | 0.09<br>(0.73) | 0.07<br>(0.73) |
| PV | -0.29<br>(0.96) | -0.16<br>(0.96) | 0.24<br>(0.96) | 0.21<br>(0.96) | 0.10<br>(0.96) | 0.03<br>(0.96) |

We have recently developed a single cell atlas combining a number of datasets to build a comprehensive atlas of the small intestine across the human life span including in utero (33). To determine if bacterially derived metabolites such as SCFA and secondary bile acids that we find in the fetal small intestine can have an in utero biological function, we explored this atlas for expression of genes associated with bile acid (*SLC10A2* (encoding ASBT), *NR1H4* (encoding FXR), *RXRA* (encoding NR2B1), *S1PR2*, *GPBAR1*, *VDR*, and *RORC*) and SCFA (*SLC5A8* (SCFA transporter), *SLC16A1* (encoding MCT1, monocarboxylate transporter), *FFAR2*, *FFAR3*, *FABP6* (encoding iBABP), and *HIF1A*) transport and signaling (**Figure 5 and Supplementary Figure S2)**. Expression of *SLC10A2*, apical sodium dependent bile acid transport carrier, significantly increased post-delivery, with minimal expression in utero. However, expression of a number of bile acid and SCFA genes was present *in utero* and restricted to subtypes of epithelial (**Figure 5A and Supplementary Figure S2A**) and or immune cells (**Figures 5B and Supplementary Figure S2B**) within the SI. The expression of *VDR* and *FABP6* steadily increased from the first trimester through adulthood with highest levels present in adults (**Figure 5C**). Nevertheless, it was still detectable in utero particularly in the second trimester with *FABP6* being expressed exclusively in the intestinal epithelial cells (IEC) within the mature absorptive (mAE) subtype and *VDR* being expressed both in the IEC (stem cells (SCs) and mAE cells) and in the immune cells (cycling Mφ, Tregs, and memory CD4 cells). Several other genes (*GPBAR1*, *SLC5A8*, *FFAR2*, *FFAR3*, *NR1H4*, *S1PR2*, and *HIF1A*) had similar expression pattern across the lifespan except for low expression in the first trimester. *FFAR2* was expressed by fetal enteroendocrine (EEC) and goblet cells as well as Mφ and ILC3s while *FFAR3* was only expressed by fetal Mφ. *NR1H4* was only expressed by IEC. *RORC* was predominantly expressed by fetal ILCs, and *HIF1A* ubiquitously expressed in all epithelia and immune cells.

**Figure 5.**
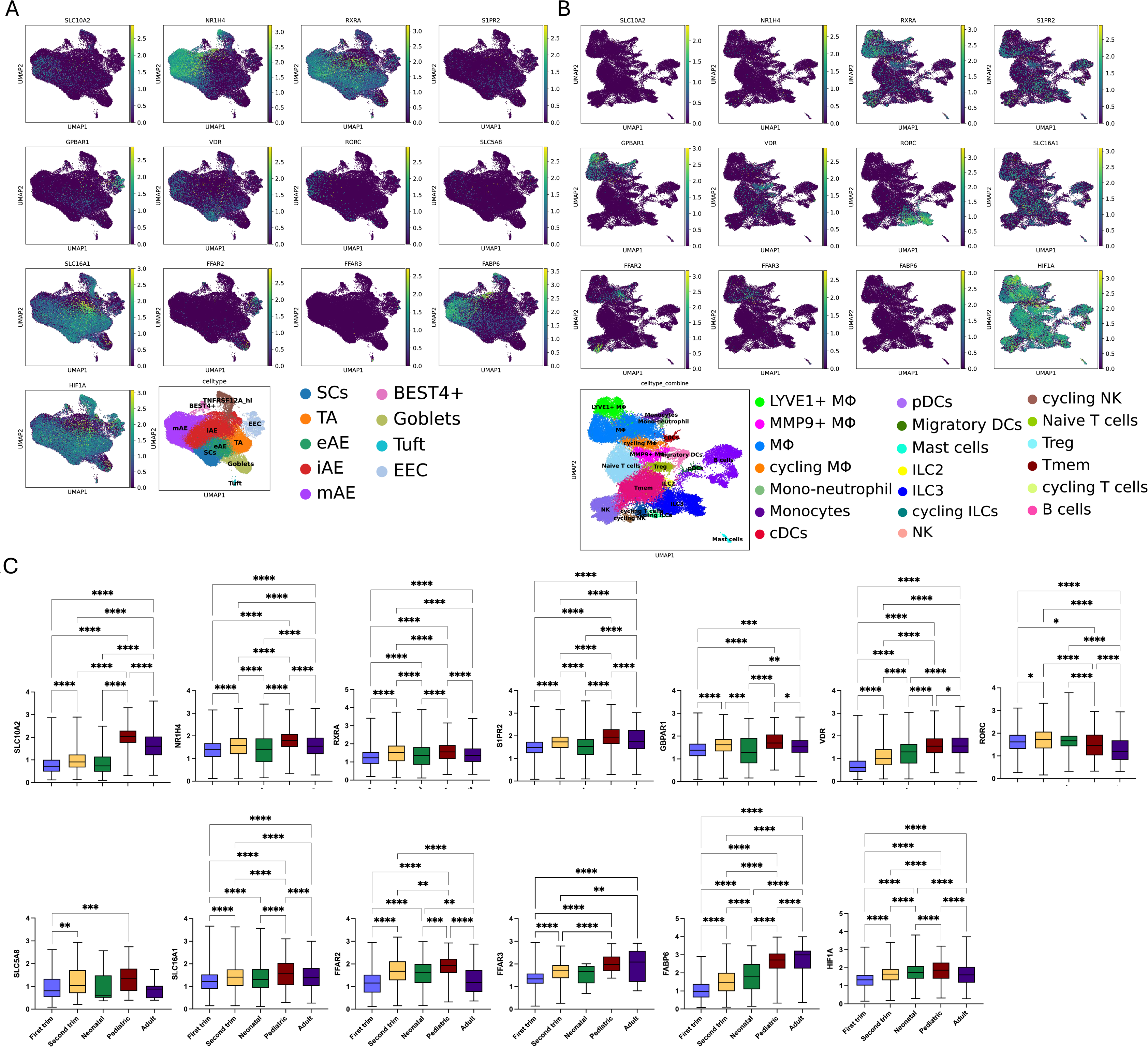
Cell type specific expression of genes associated with bile acid and SCFA transport and signaling. Uniform Manifold Approximation and Projection (UMAP) plot visualization of bile acid and SCFA associated transport and signaling gene expression in fetal small intestine epithelial (A) and fetal immune cells (B). (C) Boxplots of selected individual gene expression in small intestine across developmental stages (related to Figure 1 (33)). *P < 0.05, **P<0.01, ***P<0.001.

## Discussion

Defining when the host-microbial interactions are first established is critical for understanding normal development and identifying disease preventive strategies. Recent research has highlighted the importance of the first three years of life as critical for immune system development (4). With the recent identification of memory T cells in fetal tissues (5–10), the question arises weather intestinal T cell education and or host-microbiome interaction begins in utero, prior to delivery.

To explore in utero immune function, we had previously sought to identify potential antigens recognized by fetal intestinal T cells, but detecting in the microbiome in fetal intestines or meconium was unsuccessful (26). This is consistent with findings from other groups (12–16) who had also failed to identify microbiota in gestational tissues all suggesting that there is an absence of live microbiota in the fetal tissue. Building on murine studies (25), we previously applied the metabolomic analysis pipeline from Metabolon and provided a comprehensive report of the in utero human intestinal metabolome with an abundance of bacterially derived metabolites and aryl hydrocarbon receptor (AHR) ligands implicated in mucosal immune regulation (26). Building on these findings, in the current study, we hypothesized that microbial byproducts detected in fetal intestines in utero are primarily derived from the maternal microbiota, and travel to the human fetal intestine. To test this hypothesis, we examined a cohort of pregnancy-matched fetal tissue including the fetal intestine, the fetal meconium, the fetal placental villi, and the maternal decidua.

Consistent with our previous work, we found that the microbial metabolites were present in the fetal intestine and found that they were present in the fetus across all the tissues examined. Using our metabolomic analysis pipeline, we found that the metabolomic profile of fetal intestinal tissue was distinct from that of the fetal meconium, the fetal placental villi, and the maternal decidua, and contained bacterial metabolites. Focusing on the fetal GI tract, the top 20 most abundant metabolites contained a bacterially produced aromatic lactic acid, DHCA, that has been associated with decreasing intestinal inflammation (34–36). By Ingenuity Pathways Analysis (IPA) we further identified numerous pathways that were enriched for in the GI track compared to all other tissues that included transport of bile acids and inorganic cations, phase II conjugation of compounds and tryptophan catabolism. Tryptophan serves as the only precursor for serotonin synthesis that occurs predominantly in the intestine by EEC (37) where it plays many important functions including motility, secretion, and visceral sensitivity. When compared to meconium, additional signaling/endocrine pathways were upregulated in the fetal intestine including neurotransmitter release cycle, serotonin receptor signaling, arachidonic acid metabolism, prostaglandins and thromboxanes synthesis, and insulin secretion. We have previously demonstrated that EEC in the fetal small intestine produce insulin and all the necessary machinery for glucose sensing and insulin secretion (27). Transport of bile acids and inorganic cations is a known function of enterocytes and these data suggest that these pathways are active even in utero. This was further supported by the transcriptomics data that demonstrated that many of the bile acid transporters and receptors were expressed in utero. Phase II conjugation is a detoxifying step in drug and toxin metabolism that occur in the GI tract and liver, and this data similarly suggest that this process is active in the enterocytes in utero.

Interestingly, when compared to the meconium, many metabolic pathways were upregulated in the fetal GI tract, suggesting that fetal enterocytes are playing an active role in the absorption of amniotic fluid in utero. For example, glutamate metabolism is critical to the GI track, where glutamate has been shown to be the largest contributor to intestinal energy generation and is a precursor for glutathione, arginine and proline in the small intestine (38).

Although, live fetal microbiome does not exist in utero, maternal microbiome is important to fetal intestinal and immune development (11). Several reports suggest that neonatal mice born to mothers transiently colonized with bacteria have increased xenobiotic metabolic signatures in their intestines compared to germ-free (GF) pups and contained traces of maternal microbiome metabolites in their intestines (25). Additionally, work from the Elaine Hsiao’s group demonstrated that the maternal intestinal microbiome promotes placental development where depletion of intestinal microbiome restricted placental growth, and this was partially driven by SCFA (39). The correlation analysis performed in this study suggested that the microbial metabolites detected in fetal samples were more likely to be vertically transmitted from maternal microbiota since the fetal GI and decidua samples had substantially higher positive correlation of the abundance of the microbial metabolites than the correlation between the fetal GI and meconium samples either from matched samples or all samples analyzed together. We also investigated the variability of the abundance of the various metabolites in different tissue. Our data show that the secondary bile acid, lithocholic acid, the aromatic lactic acid, 5-MIAA, and another nine microbial metabolites were significantly enriched for in the decidua samples. Interestingly, many microbial metabolites were found in similarly abundance across all tissue samples.

Importantly, three different SCFA, butyric, isovaleric, and propionic acids, were found in fetal tissue with propionic acid being enriched for in the fetal GI tract. Butyric acid is an important energy source for enterocytes (40, 41), perhaps explaining why its levels were reduced in the GI compared to other tissue. It also plays important roles in enterocyte proliferation, differentiation and maturation (40, 41). SCFA have been shown to have anti-inflammatory roles within the intestine through binding to GPR41/43 (G protein coupled receptors, *FFAR3/2*) (42). Both *FFAR2* and *3* were expressed in the fetal intestine where *FFAR2* was found on EEC, Goblet cells, Mφ and NK cells, while *FFAR3* was found predominantly in Mφ. Secondary bile acids have also been shown to play beneficial roles in intestinal homeostasis, including regulating inflammation (43, 44). A number of genes associated with bile acid or SCFA transport and signaling were expressed in utero, especially in subtypes of epithelial and immune cells, and steadily increased from first trimester through adulthood. It’s intriguing to speculate that the maternal microbiota supports the anti-inflammatory intestinal milieu of the neonatal intestine required to induce tolerance in the setting of rapid microbiome acquisition in early life.

The main limitations of this study were the relatively small sample size, and lack of other maternal tissues including blood and stool. Though we have samples from multiple fetal tissues and maternal decidua, we were unable to secure all matched samples. Thus, the conclusions drawn about the origin of the microbial byproducts from unmatched samples may be less robust. In the future, it would be informative to collect maternal fecal and blood samples matched to placental samples for direct correlation between the maternal intestinal microbiome and bacterial metabolites in the fetal tissue.

## Methods

### Sample collection

Placental and fetal samples were obtained from the University of Pittsburgh Biospecimen Core from electively terminated products of conception (14–23 weeks’ gestation) with IRB approval and signed informed consent (IRB #18010491, University of Pittsburgh). Products of conception were collected from dilation and evacuation procedures with nonpharmacological, mechanical dilation via Dilapan-S. No fetal subjects had reported genetic abnormalities. In respect of patient confidentiality and safety, limited clinical information was collected for fetal samples. All demographic information that could be legally and respectfully obtained is shown in Table 1. After receiving fetal samples, meconium was removed, and a small piece was cut with a sterile blade and immediately snap-frozen and stored at –80°C until processing. Samples were shipped on dry ice to Khatib laboratory for metabolimics analysis.

### Un-targeted metabolomics analysis

#### Extraction method

Tissues (small and large intestinal samples) were weighed and dissolved into methanol (1:4 w/v). The samples were vortexed, homogenized, and centrifuged for 10 minutes with 15,294g at 4°C. Then the samples were filtered into HPLC vials and injected to LCMS.

#### LCMS analysis

The extracted solutions (5 μL) were injected into a UPLC connected to a photodiode array detector (Dionex Ultimate 3000), with a reverse-phase column (ZORBAX Eclipse Plus C18, 100*3.0 mm, 1.8 μm). The mobile phases consisted of phase A DDW with 0.1% formic acid and phase B acetonitrile containing 0.1% formic acid. The gradient was started with 98% A and increased to 30% B in 4 minutes, then increased to 40% B in 1 minute and kept isocratic at 40% B for another 3 minutes. The gradient increased to 50% in 6 minutes, increased to 55% in 4 minutes, and finally increased to 95% in 5 minutes and kept isocratic for 7 minutes. Phase A was returned to 98% A in 3 minutes, and the column was allowed to equilibrate at 98% A for 3 minutes before the next injection. The flow rate was 0.4 mL/min. MS analysis was performed with HESI-II source connected to a Q Exactive Plus Hybrid Quadrupole-Orbitrap Mass Spectrometer from Thermo Fisher Scientific. ESI capillary voltage was set to 3500 V, capillary temperature to 300°C, gas temperature to 350°C, and gas flow to 10 mL/min. The mass spectra (m/z 100–1500) were acquired in negative ion mode (ESI–).

Blank and quality control (QC) samples were analyzed throughout the entire experimental procedure. Blank vials consisted of methanol. The QC samples were prepared by mixing 50 μL of each sample. Blank and QC samples were injected first in the sequence, after each set of 10 samples, and at the end of the sequence, to monitor the stability and performance of the system and evaluate the quality of the acquired data.

### Metabolomic data acquisition using Compound Discoverer software

Identification, quantification and statistical analysis of peak areas identified in the various samples were executed using Compound Discoverer software (version 3.3.0.305; Thermo Scientific, Waltham, MA, USA). Molecule identification, peak determination, integration of peak area, removal of empty peaks and scale, were conducted with MZcloud (https://www.mzcloud.org) and ChemSpider (https://www.chemspider.com/) software. Results were normalized by incorporating QC samples throughout the extraction stages to check for repeatability and the extraction and normalization of the deviations. QC samples were injected during all the stages of the run to test the stability and sensitivity of the devices and to normalize these deviations.

### Quantitative metabolites analysis for validation

#### Data preprocessing

The standards and samples were injected using the same LC-MS method reported at the un-targeted metabolomics part. Peak determination and peak area integration were performed with QuanBrowser (Thermo Xcalibur, version 4.1.31.9). Autointegration was manually inspected and corrected if necessary. Calibration curves were used for the quantification of each compound. Linear curves were obtained for all compounds with R2 > 0.99: hippuric acid 0.1–5000 ppb, benzoic acid 500–50,000 ppb, taurocholate 100–50,000 ppb, and deoxycholic acid 0.1–100 ppb.

#### Method validation

Method validation was performed to determine, limit of detection (LOD), limit of quantitation (LOQ) linearity repeatability, and recovery for each compound.

For intraday precision, a mixture of all the metabolites standards was prepared and injected as QC at the beginning of the sequence, then after each 10 samples, and at the end of the sequence. The RSDs into QC samples were calculated for each analyte to be less than 6.6%.

For recovery analysis, 3 samples were spiked, extracted, and injected to LCMS. The concentration of each analyte was calculated into the spiked and nonspiked samples, and the recovery was evaluated to be on average 82% for benzoic and hippuric acids, 92% for taurocholate, and 98% for deoxycholic acid.

LOD and LOQ were determined by signal-to-noise ratios higher than 3 and 10, respectively. LOD and LOQ for hippuric acid, deoxycholic acid, and sodium taurocholate was 0.1 ppb. LOD and LOQ for benzoic acid was 500 ppb.

### Metabolomic data analysis

#### Data preprocessing

a. The original metabolome data matrix contained 65 samples and 18424 compounds. The data were preprocessed according to Li, Yujia et al. (26). The blank samples, quality control samples and NO. 55 sample without location information were filtered out, resulting in 49 samples (8 decidua, 8 small intestine, 3 large intestine, 11 small intestine meconium, 8 large intestine meconium, and 11 PV). Compounds without names were discarded, and for the named compounds with isomers, the one with the largest inter-quantile range over the 49 samples at log-scale was kept. Next, the resulting metabolome data matrix containing 49 samples and 2521 metabolites (no missing) were log-transformed (base 2) and normalized across samples by quantile normalization using the “preprocessCore” R package (45). (b) Due to sample limitation where no additional sample was available, metabolites were validated by standards however the exact concentrations of the samples could not be calculated. This raw validation metabolome data matrix contained 65 samples (same as original data matrix) and 5 compounds. After filtering out the blank samples, quality control samples, and 55 sample # 55 that failed QC, the remaining 49 samples and 5 metabolites were log-transformed (base 2). (c) Where additional samples were available, metabolites were validated by standards and exact concentrations were calculated. This raw validation data set included the quantitative abundance of 4 metabolites and 55 samples. Samples with abnormal quantity or without tissue location information were filtered out, resulting in 48 samples remaining for Hippurate, 52 samples for Benzoate, 48 samples for Taurocholic acid, and 50 samples for Deoxycholic acid. Missing values were imputed by assigning half of the minimum observed value of each metabolite and addition of a small random noise. (d) The pediatric data was from Li, Yujia et al. (26), containing 10 samples (5 large intestine and 5 small intestine) and 841 identified metabolites. According to Li, Yujia et al. (26), before log-transformation, missing values were imputed by assigning half of the minimum observed value of each metabolite and addition of a small random noise (Gaussian distribution with mean 0 and variance 100 and then rounded to the nearest integer) to the missing data to avoid the ties across samples that will impair variance estimation in subsequent differential analysis.

#### Determination of metabolite clustering

t-SNE (R package “Rtsne” (46–48)) plots were made to visualize the separability of samples from different tissue locations.

#### Statistical and bioinformatic analyses

(a) For differential analysis, the “limma” R package (49) was used to detect the differentially expressed metabolites and pathways in 3 pairwise comparisons: decidua versus GI (small intestine and large intestine), GI versus meconium, and GI versus PV. We applied “limma” R package to calculate P values and log-fold changes for individual metabolites, followed by Benjamini-Hochberg procedure to correct for multiple testing, to control false discovery rate, and to report q value. The results were visualized in volcano plots using the “EnhancedVolcano” R package (50) where the significantly differentially expressed genes are highlighted in red with the absolute value of log fold change larger than 2.5 and q value smaller than 0.05. For pathway enrichment analysis, we figured out the KEGG ID of 843 metabolites. Based on the KEGG IDs and identified differentially expressed metabolites, pathway enrichment analysis for each of the 3 pairwise comparisons were conducted using IPA software for metabolome data (QIAGEN; https://www.qiagenbioinformatics.com/products/ingenuitypathway-analysis) (51). This tool generated q values and enrichment effect sizes (log odds ratio). (c) The correlation matrix provided the pairwise correlations of the 6 tissue groups (decidua, SI, LI, SI Mec, LI Mec, and PV) from subject “1146” based on the abundance of 41 bacterially derived metabolites and was visualized by the “ggcorrplot” R package (52). The boxplots compared the correlation between GI and decidua group, and between GI and meconium group, and each point in the boxplot is the abundance correlation of microbial metabolites/Xenobiotic/fetal-derived metabolites between one decidua sample and one GI sample (or one GI sample and one meconium sample). The significance levels were calculated by two sample t-test. (d) For each microbial metabolite, bile acid and aromatic acid, we performed one-factor ANOVA and post-hoc analysis to compare the difference of its abundance among the 4 tissue groups (decidua, GI, meconium, and PV). The results were visualized in heatmap by “pheatmap” R package (53) and boxplots generated by the “ggplot2” package (54). (e) To investigate the association between the abundance of individual metabolite with gestational age and tissue group for bile acids and SCFAs, we applied the linear regression model using “limma” R package, reporting the coefficient slopes and q values.

### Single-cell RNA-seq data analysis

The single-cell data was from our previous published a single cell atlas of human small intestine throughout the human lifespan (33). The fetal epithelial cells and immune cells were clustered using Scanpy (v1.9.2) package (55) as described in Gu et al (33). Briefly, gene expression in each cell was normalized and log-transformed. Afterwards, highly variable genes were identified using the scanpy.pp.highly_variable_genes function with default parameters. In addition, the effects of the percentage of mitochondrial genes, percentage of ribosomal protein genes, and unique molecular identifier (UMI) counts were regressed out using scanpy.pp.regress_out function before scaling the data. Batch correction of samples was performed with bbknn (v1.5.1) (56). Dimensionality reduction and Leiden clustering was carried out on the remaining highly variable genes, and the cells were visualized using Uniform Manifold Approximation and Projection (UMAP) plots. Cell types were manually annotated based on known markers genes found in the literature. The statistical analysis of selected gene expression in each detected cell was performed using one-way ANOVA with Tukey’s multiple comparison test to compare gene expression among developmental stages using GraphPad Prism 9. Differences were considered statistically significant at a p value <0.05.

## Data availability

The metabolomic data is available in Github https://github.com/wenjiaking/Metabolomic-Data/tree/main, and the single-cell RNA-seq data used in this study has been listed in Supplementary Table 1 (33) where the data resources can be found.

## Author contributions

LK secured funding, and acquired all the samples. RS extracted the metabolites and performed the LC-MS experiments supervised by SK. WW analyzed the metabolomic data supervised by GT, and LK.WG performed the single-cell RNA-seq data analysis supervised by LK. OK and LK conceived of all the studies. WW and WG wrote the manuscript supervised by LK and GT. All authors have read, edited, and approved the manuscript.

## Acknowledgments

This project used the UPMC Tissue and Research Pathology/Pitt Biospecimen core which is supported in part by award P30CA047904 for fetal samples.

## Supplementary material Tables

**Supplementary Table S1. List of the detectable metabolites and their abbreviations.**

**Supplementary Table S2. Difference of metabolite abundance in pairwise comparisons between GI and the other tissue groups.**

**Supplementary Table S3. Enriched pathways in pairwise comparisons between GI and the other tissue groups**

**Supplementary Table S4. List of microbial metabolites, xenobiotics, and fetal-derived metabolites used in the analysis.**

**Supplementary Table S5. Correlation coefficients between the abundance of individual metabolites and the gestational age within each tissue group.** Except the bile acids and SCFA in Table 2, showing all the other nine bile acids and two SCFAs. The columns are the correlation coefficient within each tissue group separately. The q value for the correlation coefficient shows in parathesis *P < 0.05, **P<0.01, ***P<0.001.

## Figures

**Supplementary Figure S1. Difference validation.** Boxplots in the first row show the abundance difference between tissue groups for the eight metabolites in the main data set, while the second and third rows present the difference in validation data sets for the corresponding metabolites.

**Supplementary Figure S2. Dot plots of markers for cell type annotation.** Dot plot of the marker genes for the annotation of fetal epithelial cells (A) and of fetal immune cells (B). Color represents normalized mean expression of marker genes in each cell type, and size indicates the proportion of cells expressing marker genes.

