## Supplementary figures and images for "In utero human intestine contains maternally derived bacterial metabolites"

### Supplementary Figure S1

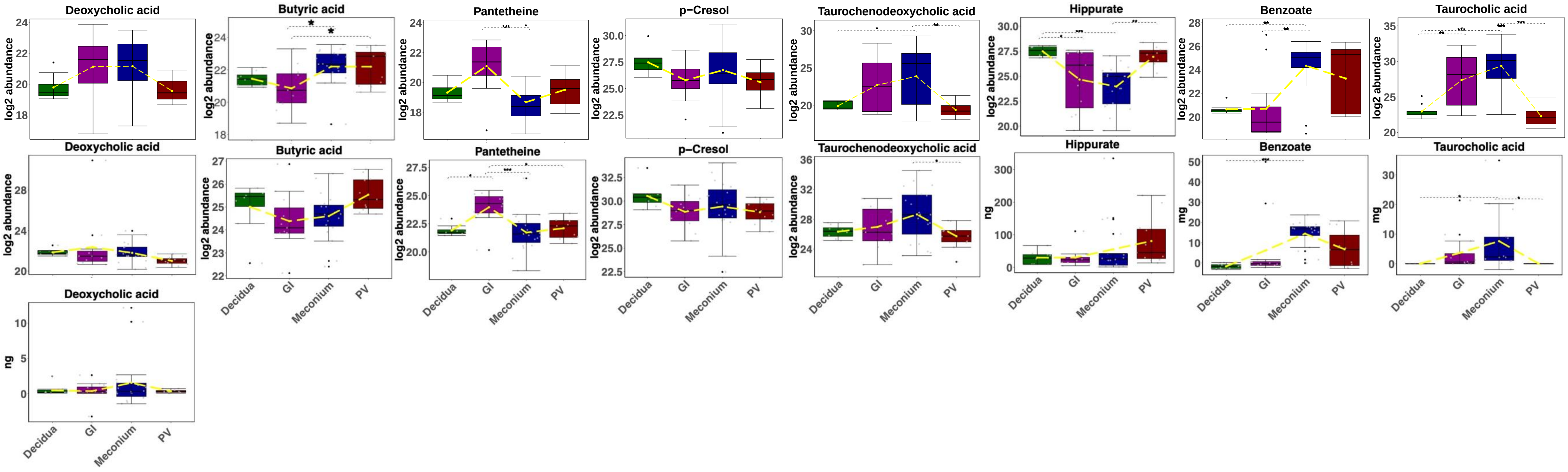

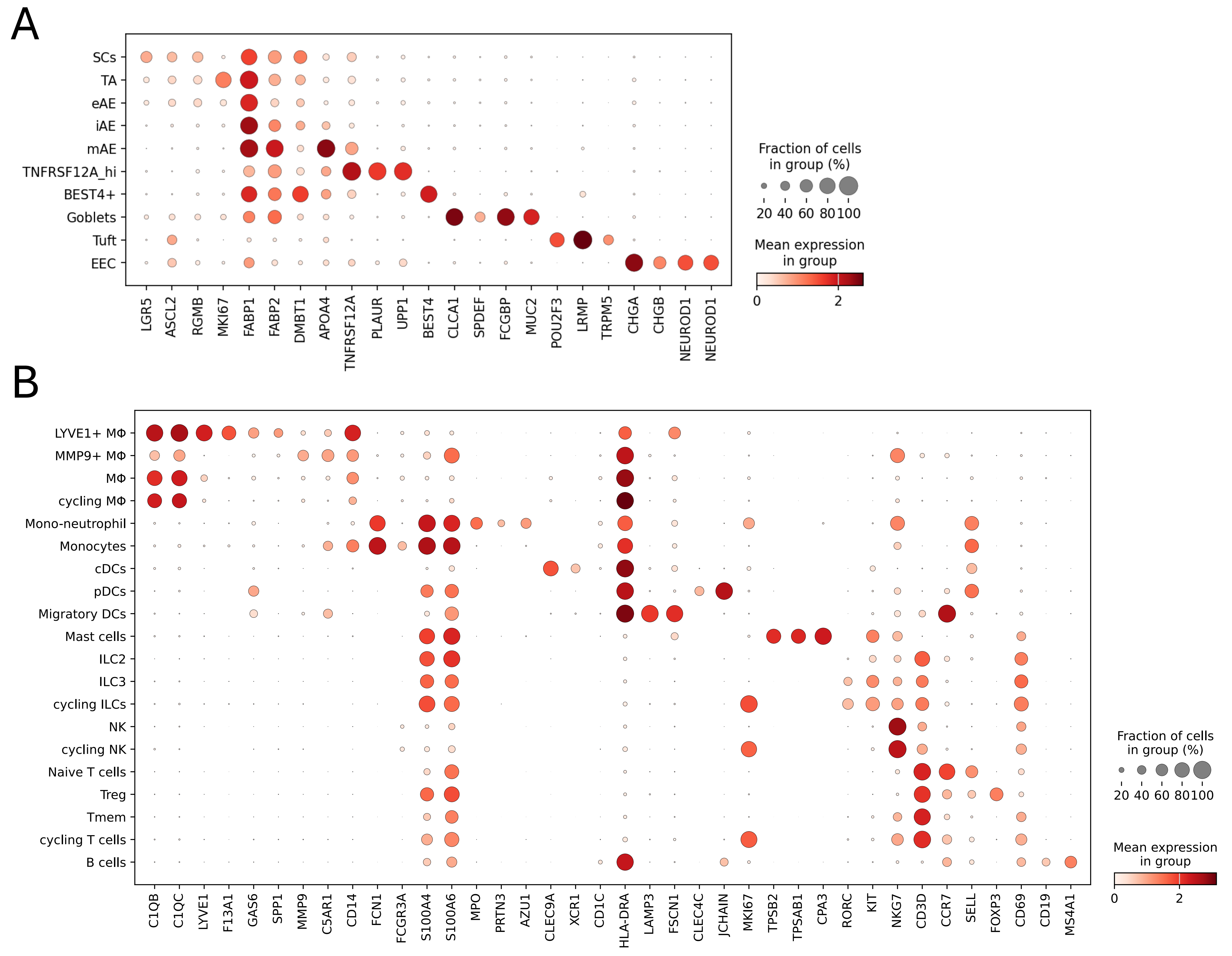

### Supplementary Figure S2

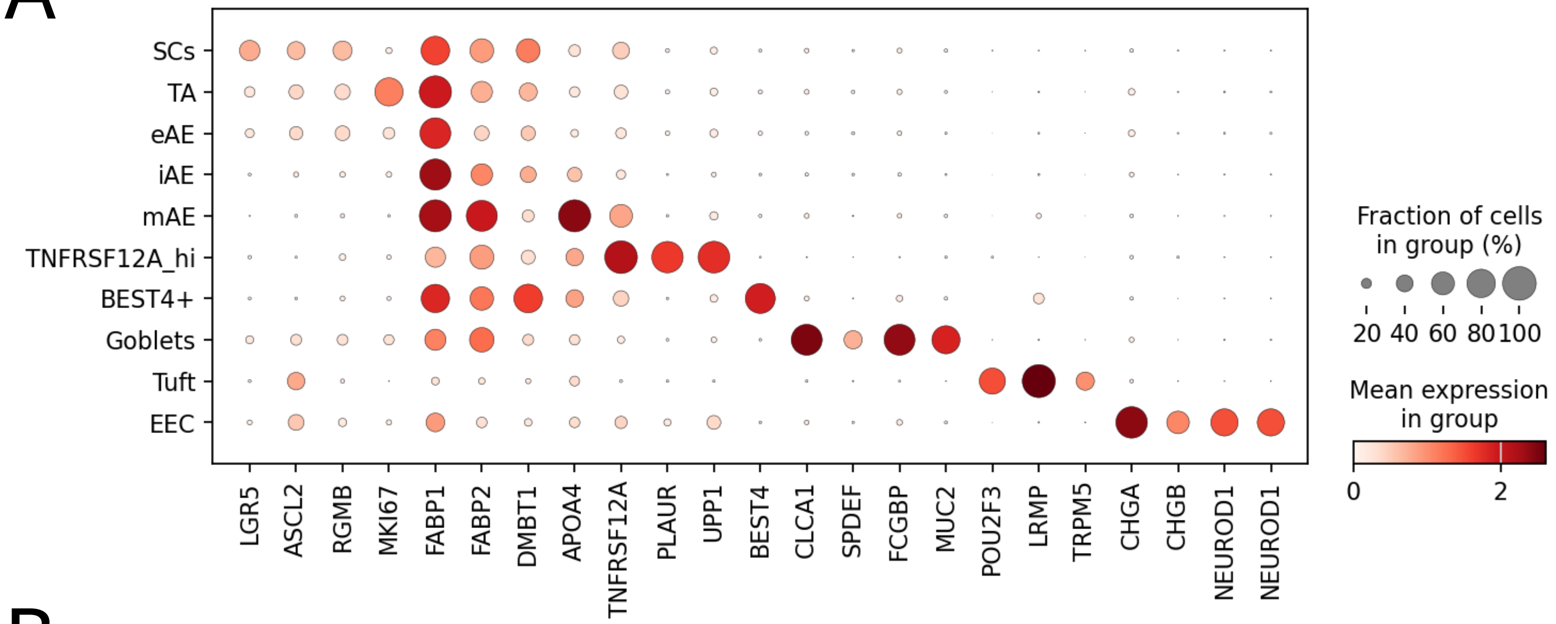

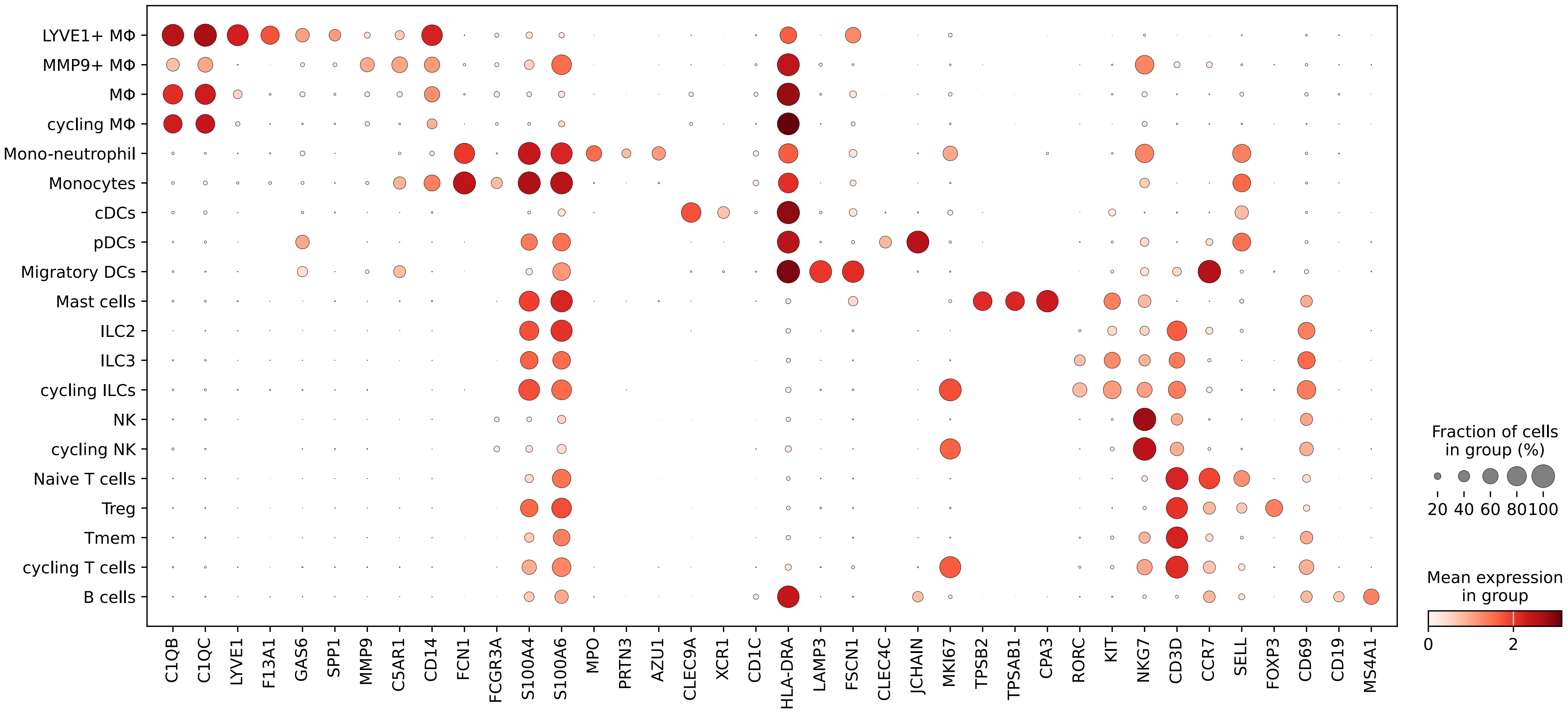
