## Supplementary Table S4 for "In utero human intestine contains maternally derived bacterial metabolites"

| Bacterial Metabolites | Xenobiotic Metabolites | Fetal-Derived Metabolites |
|---|---|---|
| Pyridoxine | Penicillin-G | N-Acetylaspartylglutamicacid |
| Pantothenic acid | 11-nor-9-Carboxy-Delta-9-THC | 5α-Pregnan-3,20-dione |
| N-Acetyl-alpha-D-glucosamine1-phosphate | (+)-Simvastatin | Pregnenolonesulfate |
| Riboflavin | (-)-Abietic acid | Homoarginine |
| Pipecolic acid | (-)-Citronellol | N-Acetylcadaverine |
| alpha-L-Arabinose | (-)-Febrifugine | Pregnenolonesulfate.1 |
| Pantetheine | (-)-Rhododendrin | 17-Hydroxypregnenolone sulfate |
| 2,2'-Methylenebis(6-tert-butyl-p-cresol) | (+)-10-Deoxymethynolide | 1beta-Hydroxycholic acid |
| Kynurenine | (+)-N-Methylconiine | Androsterone |
| Benzoate | (+/-)-Methoprene | |
| MTA | (–)-trans-Methyl dihydrojasmonate | |
| Methylhippuricacid | (1S,2R,3R,7R)-7-Isopropenyl-1-methyl-1,2,3,4,5,6,7,8-octahydro-2,3-naphthalenediol | |
| Coprocholic acid | (1S,2S,8aR)-1-[(3Z)-5-hydroxy-3-methylpent-3-en-1-yl]-2,5,5,8a-tetramethyl-decahydronaphthalen-2-ol | |
| 4-Hydroxyphenacyl alcohol | [Similar to: Nodularin; ΔMass: -623.3793 Da] | |
| albaflavenol | 18-acetoxy-1alpha-hydroxyvitamin D3 | |
| 1H-Indole | 18-acetoxy-1alpha,25-dihydroxyvitamin D3 | |
| 2-(4,6-diphenyl-1,3,5-triazin-2-yl)-5-(hexyloxy)phenol | 2-(4-Hydroxyphenyl)ethanol(Tyrosol) | |
| 4-vinylphenol sulfate | 2-(4-Nonylphenoxy)ethanol | |
| DMBS | 2-{2-[2-(Decyloxy)ethoxy]ethoxy}ethanol | |
| Phosphopantothenic acid | Asparenomycin A | |
| Phenol | Dehydrocholic acid | |
| Heppurate | Adaprolol | |
| Biotin | Alizapride | |
| Butoxyethyl phthalate | alpha-Tocotrienol | |
| Butyl-o-cresol | Ascorbyl stearate | |
| Bis(4-ethylbenzylidene)sorbitol | Cannabidiol | |
| Thiamine | Carvedilol | |
| p-Cresol | Cymoxanil | |
| 2-Hydroxyhippuric acid | Cyprodenate | |
| 5-MIAA | Doxycycline | |
| Indoxyl sulfate | Elacytarabine | |
| Deoxycholic acid | Eglumetad | |
| Glycodeoxycholic acid | Gabapentin | |
| Sulfolithocholic acid | Guaifenesin | |
| Taurodeoxycholic acid | ibufenac | |
| Lithocholic acid | Imiprothrin | |
| Butyric acid | Ipratropium | |
| Isovaleric acid | Ketamine | |
| Propionic acid | Ketorolac | |
| Dihydrocaffeic acid or DHCA | Mepivacaine | |
| Ethylparaben | Mescaline | |
| | Morphine | |
| | Nifedipine | |
| | pentobarbital | |
| | Perindopril | |
| | Picaridin | |
| | Rimexolone | |
