## Supplementary Table S5 for "In utero human intestine contains maternally derived bacterial metabolites"

| | Decidua | GI | Mecomium | PV |
|---|---|---|---|---|
| Glycochenodeoxycholic acid | 0.01<br>(0.99) | -0.17<br>(0.36) | 0.39<br>(0.43) | -0.19<br>(0.96) |
| Chenodeoxyglycocholic acid | 0.45<br>(0.94) | 0.10<br>(0.47) | -0.02<br>(0.92) | -0.02<br>(0.96) |
| Glycocholic acid | 0.51<br>(0.94) | -0.13<br>(0.46) | 0.13<br>(0.76) | 0.17<br>(0.96) |
| 7-Sulfocholic acid | 0.35<br>(0.94) | 0.19<br>(0.11) | 0.05<br>(0.92) | 0.19<br>(0.96) |
| Taurocholic acid | -0.41<br>(0.94) | -0.13<br>(0.47) | 0.33<br>(0.46) | -0.03<br>(0.96) |
| Taurodeoxycholic acid | -0.18<br>(0.94) | 0.04<br>(0.71) | 0.06<br>(0.82) | 0.02<br>(0.96) |
| Sulfolithocholic acid | 0.18<br>(0.94) | 0.25<br>(0.06) | 0.32<br>(0.43) | 0.06<br>(0.96) |
| Lithocholic acid | -0.05<br>(0.94) | 0.03<br>(0.61) | -0.06<br>(0.73) | -0.11<br>(0.96) |
| Taurochenodeoxycholic acid | -0.17<br>(0.94) | -0.09<br>(0.69) | 0.50<br>(0.43) | -0.09<br>(0.96) |
| Isovaleric acid | 0.08<br>(0.94) | 0.01<br>(0.90) | 0.02<br>(0.92) | 0.23<br>(0.96) |
| Butyric acid | -0.06<br>(0.94) | 0.00<br>(0.99) | -0.10<br>(0.73) | 0.19<br>(0.96) |
